# Diminished EBV-Specific Humoral Immunity is Associated with Neuropsychiatric Long COVID Development up to 12 Months Post-COVID-19 Symptom Onset

**DOI:** 10.64898/2026.01.27.701989

**Authors:** Philip Samaan, Patrick Budylowski, Victoria Russell, Jeevitha Srighanthan, Angela M. Cheung, Mario Ostrowski

## Abstract

Infection with SARS-CoV-2 can lead to long COVID, a chronic multisystemic condition estimated to affect approximately 400 million people worldwide. Although underlying mechanisms remain elusive, aberrant ongoing inflammation driven by Epstein-Barr virus (EBV) reactivation and persistent SARS-CoV-2 viral reservoirs have been hypothesized. We compared cellular and humoral immune responses to SARS-CoV-2 and EBV between participants with neuropsychiatric long COVID and recovered individuals. Peripheral blood mononuclear cells (PBMCs) and sera were collected from 27 long COVID individuals with ≥2 neuropsychiatric symptoms and 27 matched recovered participants at 3-6 months post-COVID-19 symptom onset (PSO). PBMCs were assessed for IFN-γ, IL-2, TNF⍺, and granzyme B T-cell responses against SARS-CoV-2, EBV, and human cytomegalovirus (HCMV). Sera were evaluated for neutralizing activity against live ancestral SARS-CoV-2 and EBV, and EBV reactivation was assessed by early antigen-diffuse IgG. We observed no significant differences in SARS-CoV-2-, EBV-, or HCMV-specific T-cell responses or live virus neutralization between long COVID and recovered groups at 3-6 months PSO. EBV reactivation was additionally only detected in one neuropsychiatric long COVID participant. However, reduced EBV neutralizing capacity at 3-6 months significantly associated with fatigue at 12 months PSO. Anti-EBV viral capsid antigen IgG levels were also significantly diminished in long COVID participants and similarly trended lower in those reporting fatigue at 12 months PSO. We therefore detected no differences in SARS-CoV-2- or EBV-specific T-cell responses or serological neutralizing capacity between neuropsychiatric long COVID and recovered participants; however, diminished EBV-specific humoral immunity may serve as a prognostic marker for neuropsychiatric long COVID development.

**IMPORTANCE:** We performed a comprehensive analysis of cellular and humoral immune responses to SARS-CoV-2, Epstein-Barr virus, and human cytomegalovirus in individuals with neuropsychiatric long COVID, a subgroup that remains poorly characterized. Although no differences in virus-specific T-cell immunity were observed between long COVID and recovered individuals, diminished Epstein-Barr virus neutralization at 3-6 months was associated with persistent or relapsing fatigue at 12 months post-COVID-19 symptom onset. Anti-viral capsid antigen IgG antibody levels were also significantly lower in neuropsychiatric long COVID participants at 3-6 months and similarly trended lower in those reporting fatigue at 12 months post-symptom onset. Together, these findings suggest that impaired humoral immunity to Epstein-Barr virus may contribute to the development or persistence of neuropsychiatric long COVID and highlight a promising future direction for mechanistic studies.

## INTRODUCTION

Per the definition developed by the National Academies of Sciences, Engineering, and Medicine (NASEM) in July 2024 (1), post-COVID-19 condition (long COVID) “*…is an infection-associated chronic condition (IACC) that occurs after SARS-CoV-2 infection and is present for at least 3 months as a continuous, relapsing and remitting, or progressive disease state that affects one or more organ systems.”* Globally, it is estimated that 36% of infected individuals experience long COVID (2), with approximately 400 million cases worldwide (3). Due to its multisystemic effects, long COVID may be reported as a complex mixture of respiratory, gastrointestinal, cardiovascular, and neuropsychiatric symptoms (4).

Despite its prevalence, the pathogenesis of long COVID remains poorly understood. Emerging evidence suggests that chronic immune activation – potentially driven by a persistent SARS-CoV-2 reservoir and the reactivation of latent herpesviruses – may contribute to ongoing symptomatology (5–14). Studies detecting SARS-CoV-2 spike and nucleoprotein antigens in the plasma and lung tissue of 65-70% of long COVID participants several months after infection (10, 11) parallel reports of sustained innate and adaptive immune activation in long COVID compared with asymptomatic recovered individuals at similar timepoints (5, 6). B-cell activation during the antiviral immune response may trigger the reactivation of Epstein-Barr virus (EBV), a gammaherpesvirus that establishes lifelong latent B-cell reservoirs in over 95% of the global population (15). In a cohort studied by Gold et al., 67% of individuals reporting long COVID symptoms ≥90 days post-infection exhibited serological evidence of EBV reactivation compared with only 10% of recovered participants, and anti-early antigen diffuse (EA-D) IgG antibody titers positively correlated with symptom burden (12). Similarly, Rohrhofer et al. observed that long COVID individuals experiencing fatigue at a median of 235 days post-infection were more than twice as likely to display DNA evidence of EBV reactivation in throat washings relative to recovered individuals (14). As such, current evidence collectively suggests that chronic inflammation driven by persistent viral antigen or secondary EBV reactivation may underlie long COVID pathogenesis.

Effective T- and B-cell responses are critical for SARS-CoV-2 clearance (16–25) and maintaining lifelong EBV control (26–28). However, studies examining SARS-CoV-2- and EBV-specific T- or B-cell immunity in long COVID are highly diverse and altogether inconclusive. Further characterization of these immune responses is thus highly warranted for understanding disease pathogenesis.

The cellular response to an acute SARS-CoV-2 infection is dominated by a CD4^+^ T-cell response characterized by the Th1 cytokines IFN-𝛾, TNF, and IL-2 (29–31). Although these cytokines are proposed to bolster the immune response to viral infections, SARS-CoV-2 may paradoxically exploit their systemic effects to enhance viral infection and replication within tissues, leading to a prolonged disease course in some individuals. Previous studies using *in vitro* organoid models have reported that IFN-𝛾 and TNF can upregulate ACE2 expression on human enterocytes (32) and cardiomyocytes (33), respectively, leading to enhanced viral infection and replication. Furthermore, in chronic viral infections, persistent IL-2 signaling can drive CD8^+^ T-cell exhaustion (34). Multiple studies have already reported on the elevated exhaustion and diminished responsiveness of SARS-CoV-2-specific CD8^+^ T-cells in long COVID compared with recovered individuals (35–37). Given the increasing evidence of a persistent SARS-CoV-2 reservoir (7–11), we thus hypothesized that long COVID individuals would exhibit elevated Th1-type (IFN-𝛾, TNF, and IL-2) and diminished cytotoxic (granzyme B) T-cell responses to SARS-CoV-2 compared to recovered individuals. Since pro-inflammatory T-cell signaling can also induce B-cell activation and thereby promote the reactivation of latent EBV reservoirs (38), we further hypothesized that EBV-specific T-cell responses would be elevated in long COVID participants.

B-cell dysregulation may likewise contribute to long COVID pathogenesis. Regarding SARS-CoV-2, delayed neutralizing antibody responses have been linked to fatal COVID-19 outcomes (25), and long COVID individuals reportedly exhibit significant reductions in SARS-CoV-2-specific humoral immunity several months following infection compared with recovered individuals (22–24). With respect to EBV, deficiencies in viral capsid antigen (VCA)- and Epstein-Barr nuclear antigen 1 (EBNA1)-specific memory B-cells have been observed in individuals with chronic fatigue syndrome and correlated with enhanced latent EBV replication in infected B-cells (28). Immunodeficient mouse models reconstituted with human cord blood-derived CD34^+^ hematopoietic stem cells have additionally demonstrated that EBV-neutralizing antibodies are essential for limiting viral replication and preventing downstream EBV-associated malignancies, including lymphoproliferative disorders and cancer, following primary infection (39, 40). We therefore hypothesized that SARS-CoV-2- and EBV-specific neutralizing antibody titres would be diminished in long COVID compared to recovered individuals.

Investigating the immunological mechanisms of long COVID is essential for identifying biomarkers of disease pathogenesis that can inform the development of novel targeted therapeutic strategies aimed at improving clinical outcomes and mitigating the long-term impacts of COVID-19. To pursue our hypotheses, four-colour FluoroSpot assays were deployed to assess peripheral blood mononuclear cell (PBMC) samples from long COVID and recovered individuals for IL-2, IFN-γ, TNF, and GzmB T-cell responses against SARS-CoV-2 and EBV at 3-6 months post-COVID-19 symptom onset (PSO). HCMV was additionally included in our analysis to investigate a previous report of a negative association between long COVID incidence and HCMV seropositivity (41). Live virus neutralization assays were further conducted to evaluate sera for neutralizing capacity against SARS-CoV-2 and EBV.

## METHODS

### Sample Acquisition

Cryopreserved PBMC and serum samples were acquired from long COVID and recovered participants enrolled in the Canadian COVID-19 longitudinal cohort study (CANCOV; ClinicalTrials.gov ID NCT05125510). Samples were retrieved from University Health Network Biobank, Toronto, Ontario, Canada in compliance with the Declaration of Helsinki (CTO 2157).

### Assessing SARS-CoV-2- and EBV-Specific T-cell Responses by FluoroSpot

Four-colour FluoroSpot Flex kits from MabTech were used in accordance with the manufacturer’s instructions to interrogate PBMC samples for IFN-γ, TNF, IL-2, and granzyme B (GzmB) T-cell responses to SARS-CoV-2 (nucleoprotein [102 peptides, JPT] and spike [315 peptides, JPT]) and EBV (BZLF1 [59 peptides, JPT] and EBNA-1 [158 peptides, JPT]) protein masterpools. The masterpools were comprised of 15mer peptides with 11 amino acid (a.a.) overlaps spanning the entirety of their respective antigens and used at a concentration of 1 ug/mL to stimulate PBMCs for a period of 48 hours at 37°C, 5% CO_2_.

As a positive control to assess T-cell viability and functionality, a masterpool comprised of 138 15mer peptides with 11a.a. overlaps spanning the HCMV tegument protein – pp65 (NIH AIDS Reagents Program) – was used at a concentration of 1 ug/mL. Staphylococcal enterotoxin B (SEB) was used at a concentration of 0.1 ug/mL as a positive control for the assay’s functionality. PBMCs cultured in medium only with 0.2% (v/v) DMSO – the final DMSO concentration in all treatment wells – were used as negative controls.

FluoroSpot readings were conducted using the MabTech IRIS II^TM^ analyzer. To analyze the data, the sum of spots across duplicate negative control wells was subtracted from the sum of spots across duplicate treatment wells.

### Live Ancestral SARS-CoV-2 Neutralization Assays

All live ancestral SARS-CoV-2 neutralization assays were conducted using Vero E6 cells as a substrate, as previously described (42).

### Anti-VCA IgG ELISA Assay

Anti-VCA IgG antibody levels were quantified in serum samples using the SERION ELISA classic EBV VCA IgG kit (Cat# ESR1361G, QED Bioscience, CA, USA), as per the manufacturer’s instructions.

### Anti-EA-D IgG ELISA Assay

Anti-EA-D IgG antibody levels were quantified in serum samples using the Creative Diagnostics EA-D IgG ELISA kit (Cat# DEIA329, Creative Diagnostics, NY, USA), as per the manufacturer’s instructions.

### Anti-HCMV IgG ELISA Assay

Anti-HCMV IgG antibody levels were semi-quantitatively measured in serum samples using the Abnova Cytomegalovirus IgG ELISA kit (Cat# KA1452, Abnova, Taipei, Taiwan), as per the manufacturer’s instructions.

### Cell Lines

HONE-Akata-Z cells are HONE-1 Akata epithelial cells containing EBV that expresses GFP as well as an integrated doxycycline-inducible BZLF1 for lytic cycle induction (43). These cells were cultured in Alpha-MEM medium supplemented with 10% heat-inactivated fetal bovine serum (FBS; Wisent), 10mM HEPES (Wisent), and 100UI/mL penicillin-streptomycin (Wisent). EBV-negative Akata B-cells were cultured in R10 medium. This consisted of RPMI 1640 medium (Wisent) supplemented with 10% heat-inactivated FBS, 10 mM HEPES, 2 mM GlutaMAX (Thermo Fisher Scientific), 100 UI/mL of penicillin-streptomycin, and 1 mM sodium pyruvate (Thermo Fisher Scientific). Both cell lines were generously donated by Dr. Lori Frappier’s laboratory (University of Toronto).

### Induction, Harvesting, and Concentration of GFP-Expressing EBV

HONE-Akata-Z cells were grown to 80% confluency, then treated with 2ug/mL of doxycycline (Millipore Sigma, D9891) to induce lytic reactivation of GFP-expressing EBV. At 72-hours post-induction, cell culture supernatant containing the virus was transferred to a 50mL Falcon tube and centrifuged for 5 minutes at 500xg to remove cell debris. To further purify and concentrate the viral isolate, the supernatant was passed through a 0.45um PE syringe filter (FroggaBio, SF0.45PES) into a 10kDa Amicon-15 Ultracentrifuge tube (Millipore Sigma, UFC9010) and centrifuged at 5,000xg for 30 minutes at 4°C. The concentrated viral supernatant was then aspirated and aliquoted for storage at −80°C until needed.

### TCID_50_ Assay for Titrating GFP-Expressing EBV

The concentrated stock of GFP-expressing EBV was titrated using the TCID_50_ assay. Briefly, 10x serial dilutions of stock were created in replicates of 12 from rows A to H of a 96-well round bottom polystyrene plate (dilutions ranged from 10^−1^ to 10^−8^). EBV-negative Akata B-cells were seeded at 25,000 cells per well and incubated for 3 hours at 37°C, 5% CO_2_. The plate was then centrifuged at 500xg for 5 minutes before the viral inoculum was discarded and replaced with fresh R10 medium. Following a 48-hour incubation period at 37°C, 5% CO_2_, the plate was washed with 2% heat-inactivated FBS in D-PBS and fixed with 4.2% paraformaldehyde buffer (BD Biosciences, 554655) in preparation for reading by a BD LSRFortessa X-20 Cell Analyzer. EBV-infected B-cells fluoresced with GFP and were clearly detectable within the Blue 525/50 channel. Using a positivity threshold of 0.1% infection per well, the TCID_50_ of the virus was calculated using the Reed-Muench method.

### Live EBV Neutralization Assay

Serum samples were first incubated at 56°C for 30 minutes to inactivate complement. A 10x serial dilution of each sample was then performed in duplicate from columns 1 to 11 of a 96-well round bottom plate using RPMI 1640 medium as a diluent. Samples were co-cultured with 200 TCID_50_ of GFP-expressing EBV and incubated for 1 hour at 37°C, 5% CO_2_. The twelfth column of the plate was reserved as a virus-only positive control. Following incubation, 25,000 EBV-negative Akata B-cells were seeded per well and incubated for 3 hours at 37°C, 5% CO_2_. The serum/virus co-culture was subsequently removed from all wells and replaced with fresh R10 medium prior to a final incubation period of 48 hours at 37°C, 5% CO_2_. Following incubation, the cells were fixed using 4.2% paraformaldehyde buffer and EBV infectivity was read the same day using the Blue 525/50 channel of the BD LSRFortessa X-20 Cell Analyzer. %Neutralization (%NT) in each well was calculated as follows:

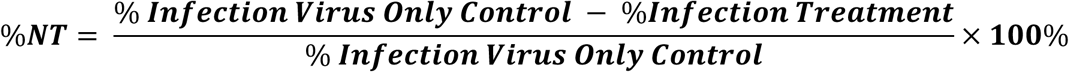

A 4-parameter non-linear regression analysis was performed using Prism GraphPad software (GraphPad Software, La Jolla, CA) to compute the 50% inhibitory concentration (IC_50_) of each serum sample assayed.

### Fluorescence Microscopy Image Enhancement

A fluorescence microscopy image of EBV-infected Akata B-cells in this report (**Figure 4A**) was enhanced by adjusting the gamma signal using Adobe Photoshop 2026.

### Statistical Analysis

All statistical analysis was conducted using Prism GraphPad software. Cross-sectional and correlational analyses were performed using two-tailed Mann-Whitney U tests and Spearman correlation tests, respectively. P-values in **Table 1** were derived using the Mann-Whitney U test or Fisher’s exact test, as appropriate. * = 0.01 < P ≤ 0.05, ** = 0.001 < P ≤ 0.01, *** = 0.0001 < P ≤ 0.001, **** = P ≤ 0.0001.

**Table 1:** Clinical characteristics of CANCOV study participants.

|  |  | Long COVID<br>(n = 27) | Recovered<br>(n = 27) | Total<br>(n = 54) | p-value |
| --- | --- | --- | --- | --- | --- |
| Sex | Female | 19 (70.4%) | 17 (63.0%) | 36 (67.0%) | 0.7734 |
|  | Male | 8 (29.6%) | 10 (37.0%) | 18 (33.0%) |  |
| Age | Median | 46.0 | 43.0 | 45.5 | 0.8456 |
|  | IQR | 16.0 | 20.0 | 16.0 |  |
| Age Group | 20-34 | 4 (14.8%) | 6 (22.2%) | 10 (18.5%) | 0.5164 |
|  | 35-49 | 11 (40.7%) | 11 (40.7%) | 22 (40.7%) |  |
|  | 50-64 | 11 (40.7%) | 7 (25.9%) | 18 (33.3%) |  |
|  | 65-79 | 1 (3.7%) | 3 (11.1%) | 4 (7.4%) |  |
| BMI | Median | 25.1 | 26.2 | 25.8 | 0.9316 |
|  | IQR | 6.31 | 4.27 | 5.98 |  |
| Ethnicity | Caucasian | 19 (70.4%) | 16 (59.3%) | 35 (64.8%) | 0.7985 |
|  | Latin American | 2 (7.4%) | 1 (3.7%) | 3 (5.6%) |  |
|  | East Asian | 1 (3.7%) | 2 (7.4%) | 3 (5.6%) |  |
|  | South Asian | 1 (3.7%) | 0 (0.0%) | 1 (1.9%) |  |
|  | Black African | 0 (0.0%) | 1 (3.7%) | 1 (1.9%) |  |
|  | Black Caribbean | 1 (3.7%) | 3 (11.1%) | 4 (7.4%) |  |
|  | Middle Eastern | 1 (3.7%) | 3 (11.1%) | 4 (7.4%) |  |
|  | First Nations | 1 (3.7%) | 0 (0.0%) | 1 (1.9%) |  |
|  | Mixed Heritage | 1 (3.7%) | 1 (3.7%) | 2 (3.7%) |  |
| SARS-CoV-2 Wave<sup>1</sup> | 1 | 1 (3.7%) | 0 (0.0%) | 1 (1.9%) | 0.5040 |
|  | 2 | 10 (37.0%) | 14 (51.9%) | 24 (44.4%) |  |
|  | 3 | 11 (40.7%) | 6 (22.2%) | 17 (31.5%) |  |
|  | 4 | 1 (3.7%) | 2 (7.4%) | 3 (5.6%) |  |
|  | 5 | 4 (14.8%) | 5 (18.5%) | 9 (16.7%) |  |
| Months PSO | 3 | 3 (11.1%) | 6 (22.2%) | 9 (16.6%) | 0.4577 |
|  | 4 | 3 (11.1%) | 3 (11.1%) | 6 (11.1%) |  |
|  | 5 | 2 (7.4%) | 0 (0.0%) | 2 (3.7%) |  |
|  | 6 | 19 (70.4%) | 18 (66.7%) | 37 (68.5%) |  |

<sup>1</sup> **Wave 1:** Ancestral (March 2020); **Wave 2:** Ancestral (September 2020 – February 2021); **Wave 3:** Ancestral + Alpha (March – May 2021); **Wave 4:** Delta (September – December 2021); **Wave 5:** Delta + BA.1 (December 2021 – January 2022)

|  |  |  |  |  |  |
| --- | --- | --- | --- | --- | --- |
| # of SARS-CoV-2 Vaccine Doses Received | 0 | 4 (14.8%) | 9 (33.3%) | 13 (24.1%) | 0.3882 |
|  | 1 | 5 (18.5%) | 4 (14.8%) | 9 (16.7%) |  |
|  | 2 | 16 (59.3%) | 11 (40.7%) | 27 (50.0%) |  |
|  | 3 | 2 (7.4%) | 3 (11.1%) | 5 (9.3%) |  |
| HCMV Positivity (Anti-HCMV IgG) | Negative | 13 (48.1%) | 13 (48.1%) | 26 (48.1%) | >0.9999 |
|  | Positive | 14 (51.9%) | 14 (51.9%) | 28 (51.9%) |  |
| EBV Positivity (Anti-VCA IgG) | Negative | 2 (7.4%) | 4 (14.8%) | 6 (11.1%) | 0.6687 |
|  | Positive | 25 (92.6%) | 23 (85.2%) | 48 (88.9%) |  |
| EBV Reactivation (Anti-EA-D IgG) | Negative | 26 (96.3%) | 27 (100%) | 53 (98.1%) | >0.9999 |
|  | Positive | 1 (3.7%) | 0 (0.0%) | 1 (1.9%) |  |

### Data Included within Manuscript

All data supporting the findings of this study are provided within the article.

## RESULTS

### Clinical Characteristics

A total of 27 long COVID and 27 recovered participants – matched for age, sex, BMI, ethnicity, and time of sampling – were enrolled into our study from the CANCOV cohort. CANCOV serves as a longitudinal observational study platform that aims to evaluate the physical and mental health impacts of COVID-19 on participants and their caregivers up to 5 years from initial diagnosis. The cohort includes approximately 2000 adult participants across Ontario, Quebec, British Columbia, Alberta, and Manitoba (ClinicalTrials.gov ID NCT05125510). All participants included in our study were non-hospitalized with an initially mild symptomatic SARS-CoV-2 infection. Clinical characteristics can be found in **Table 1**. Briefly, our cohort was predominantly female (67.0%) with a median age of 45.5 years (IQR = 16.0) and a median BMI of 25.8 (IQR = 5.98). The majority of individuals were Caucasian (64.8%), infected within the second (44.4%) or third (31.5%) waves of the pandemic – corresponding to the ancestral and alpha waves from September 2020 to May 2021 (44) – and sampled at 6 months PSO (68.5%). Long COVID and recovered participants were also matched for the number of vaccine doses received prior to the time of sampling, with the majority receiving 2 vaccine doses (50.0%). With respect to herpesvirus infections, the two groups were well matched for EBV and HCMV seropositivity.

Under the 2024 NASEM criteria, individuals presenting with at least one symptom for ≥3 months following SARS-CoV-2 infection may be clinically diagnosed with long COVID (1). However, the application of these diagnostic criteria to COVID-19-negative individuals has been shown to yield a high false-positive rate of 28-40% at 3 months and 15-23% at 6 months post-acute illness, reflecting a considerable background prevalence of commonly reported persistent symptoms among the general population (45). Neuropsychiatric symptoms are the most frequently reported among long COVID individuals (46). Thus, to improve the specificity of our long COVID classifications and minimize confounding effects in our analyses, we restricted our focus to long COVID participants who exhibited ≥2 neuropsychiatric symptoms at 3-6 months PSO. These symptoms included fatigue, cognitive deficits, change in taste, change in smell, numbness/tingling, tinnitus, headache, vertigo, dizziness, seizures, sudden loss of consciousness, insomnia, anxiety, and depression. Groups in our analysis could thus be defined as follows:

1. **Long COVID (LC) group:** Initially symptomatic and not hospitalized during a mild SARS-CoV-2 infection with ≥2 ongoing or relapsing neuropsychiatric symptoms (fatigue, cognitive dysfunction, etc.) reported at 3-6-months PSO; and
2. **Recovered (R) group:** Initially symptomatic and not hospitalized during a mild SARS-CoV-2 infection with ≤1 reported symptom at 3-6-months or 12-months PSO.

As shown in **Figure 1A-C**, long COVID individuals reported up to 29 symptoms at 3-6 months PSO and up to 30 symptoms at 12 months PSO. Fatigue was predominantly reported in 52-56% of participants across all study timepoints, while cognitive deficits were also commonly reported in 56% and 48% of participants at 6 and 12 months PSO, respectively.

**Figure 1:**
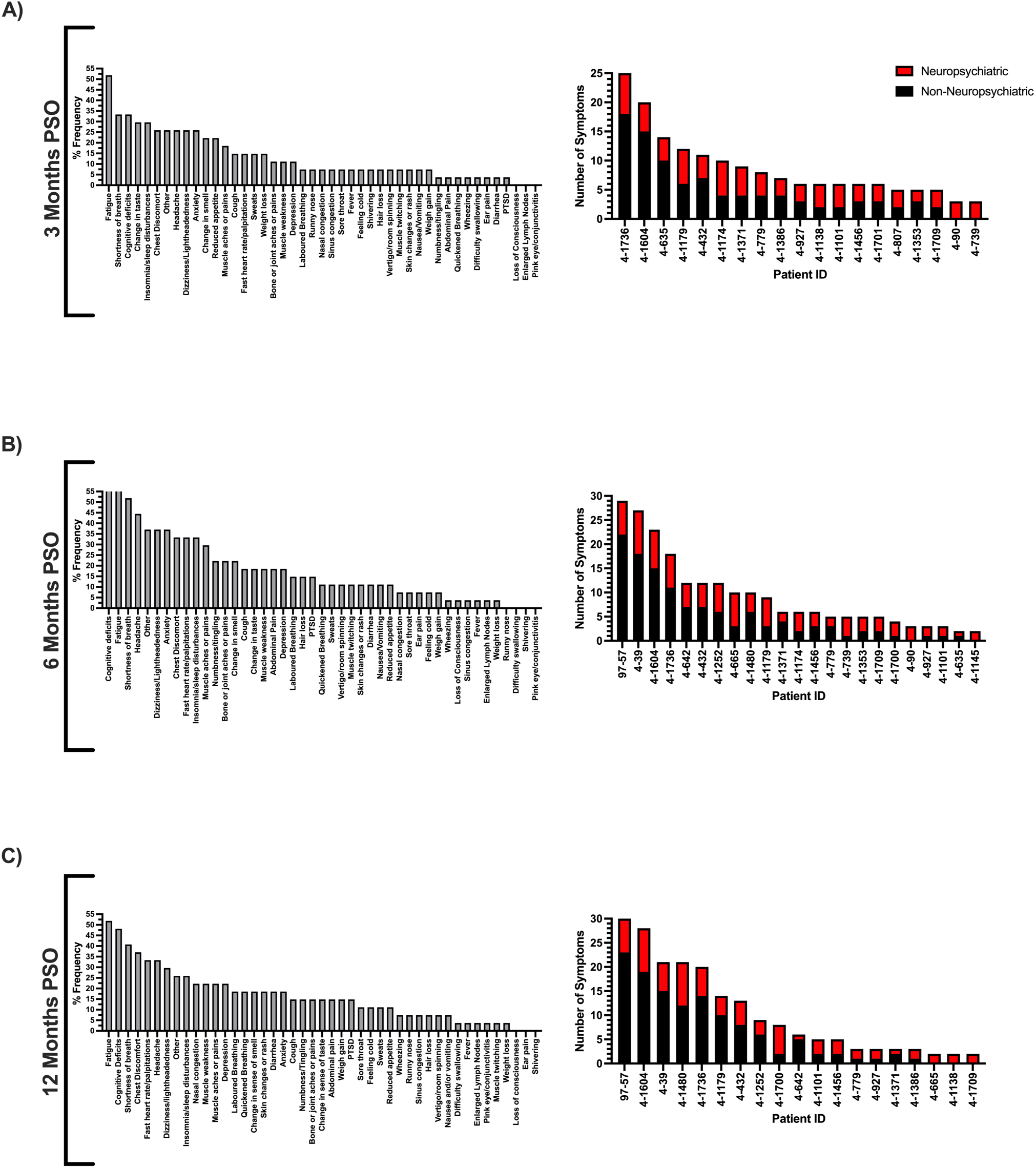
Percentage frequencies and number of symptoms experienced among long COVID participants at (A) 3 months, (B) 6 months, and (C) 12 months PSO.

Within our recovered group, 4 participants reported one symptom of either mild fatigue or mild-moderate chest discomfort at 3-6 months PSO. The remaining 23 reported no symptoms at 3-6 months or 12 months PSO.

### A Comparison of SARS-CoV-2-, EBV-, and HCMV-Specific IFN-**γ,** IL-2, TNF, and GzmB T-cell Responses between Neuropsychiatric Long COVID and Recovered Participants at 3-6 Months PSO

To pursue our hypothesis regarding long COVID T-cell responses against SARS-CoV-2 and EBV, 4-colour FluoroSpot assays were conducted *ex vivo* on PBMCs to measure virus-specific effector cytokine responses. For SARS-CoV-2, IFN-γ, IL-2, TNF, and GzmB T-cell responses were measured in response to nucleoprotein (N) and spike (S) peptide masterpools. For EBV, the same effector cytokine responses were measured in response to EBNA1 and BZLF1 peptide masterpools. The HCMV tegument protein – pp65 – was also included in our analysis to evaluate a prior report of a negative correlation between HCMV seropositivity and long COVID incidence (41). T-cell responses against EBV and HCMV were only assessed in individuals who were seropositive for anti-VCA IgG or anti-HCMV IgG, respectively.

In light of the considerable degree of variability within groups, no significant differences in SARS-CoV-2-, EBV-, or HCMV-specific T-cell responses could be detected between neuropsychiatric long COVID and recovered participants (**Figure 2**).

**Figure 2:**
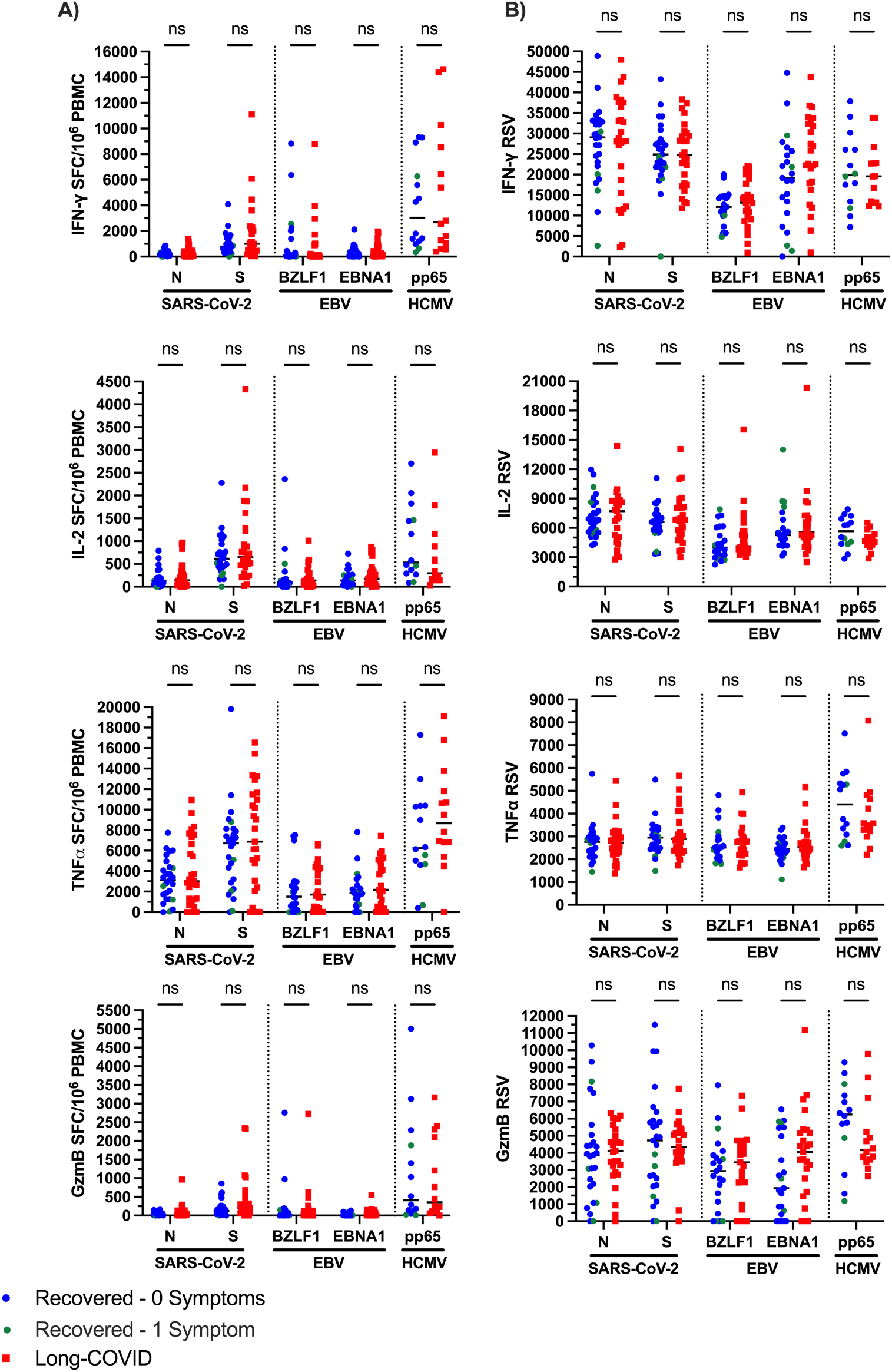
A comparison of IFN-γ, IL-2, TNF, and GzmB T-cell responses between neuropsychiatric long COVID and recovered participants. (A) Spot forming cells (SFC) per 10^6^ PBMC and (B) relative spot volumes (RSV) were quantified against SARS-CoV-2 (N, S), EBV (BZLF1, EBNA1), and HCMV (pp65). Horizontal solid lines represent medians.

### A Comparison of Neutralizing Antibody Levels against Live Ancestral SARS-CoV-2 between Neuropsychiatric Long COVID and Recovered Participants at 3-6 Months PSO

Live SARS-CoV-2 neutralization assays were performed on participant sera obtained at 3-6 months PSO using Vero E6 cells as a substrate. As shown in **Figure 3**, no differences in SARS-CoV-2 neutralizing capacity could be observed between neuropsychiatric long COVID and recovered participants.

**Figure 3:**
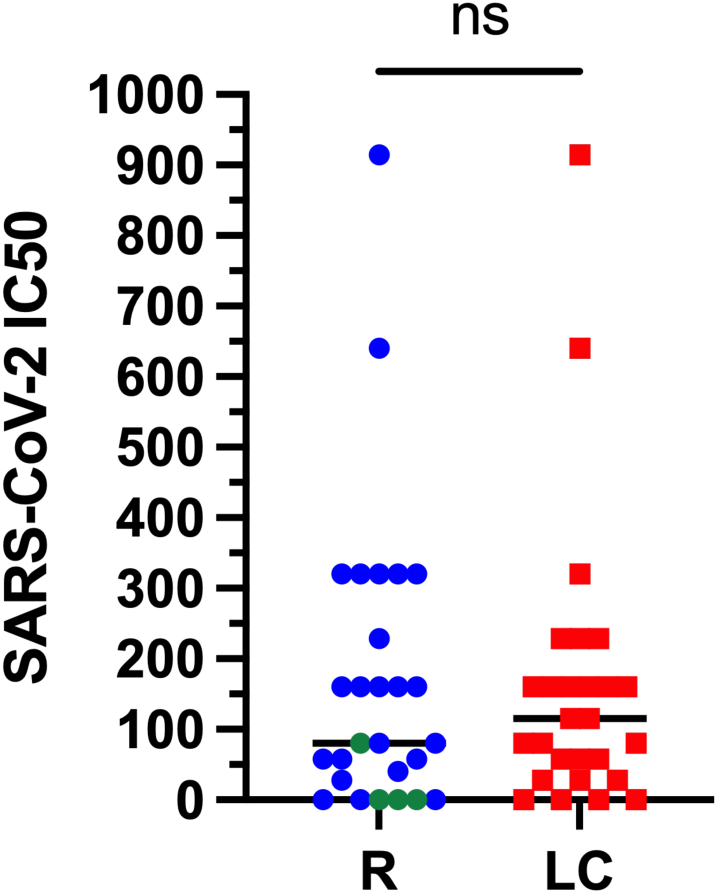
A comparison of live ancestral SARS-CoV-2 neutralizing antibody levels in neuropsychiatric long COVID and recovered participants. Horizontal solid lines represent median values. R = recovered (blue circles = 0 symptoms; green circles = 1 symptom); LC = long COVID.

### Assessing EBV Neutralizing Capacity and Reactivation in Long COVID and Recovered Participants at 3-6 Months PSO

Traditionally, EBV neutralizing antibody titers are measured using B-cell transformation assays, which assess the ability of EBV to transform primary B-cell cultures in the presence of human plasma or serum. However, this approach is often labor-intensive and requires 4-6 weeks to perform. Accordingly, a more sensitive and high throughput method involving the use of GFP-expressing EBV has been adopted from Li et al. (47) and modified in our work to accurately assess EBV-neutralizing antibody titers within human plasma or serum by flow cytometry (see Methods). Similarly to Li et al. (47), we have shown that the Akata EBV-negative B-cell line is highly susceptible to infection with the EBV-GFP virus, deeming it as an appropriate substrate for our live EBV neutralization assays (**Figure 4A**).

**Figure 4:**
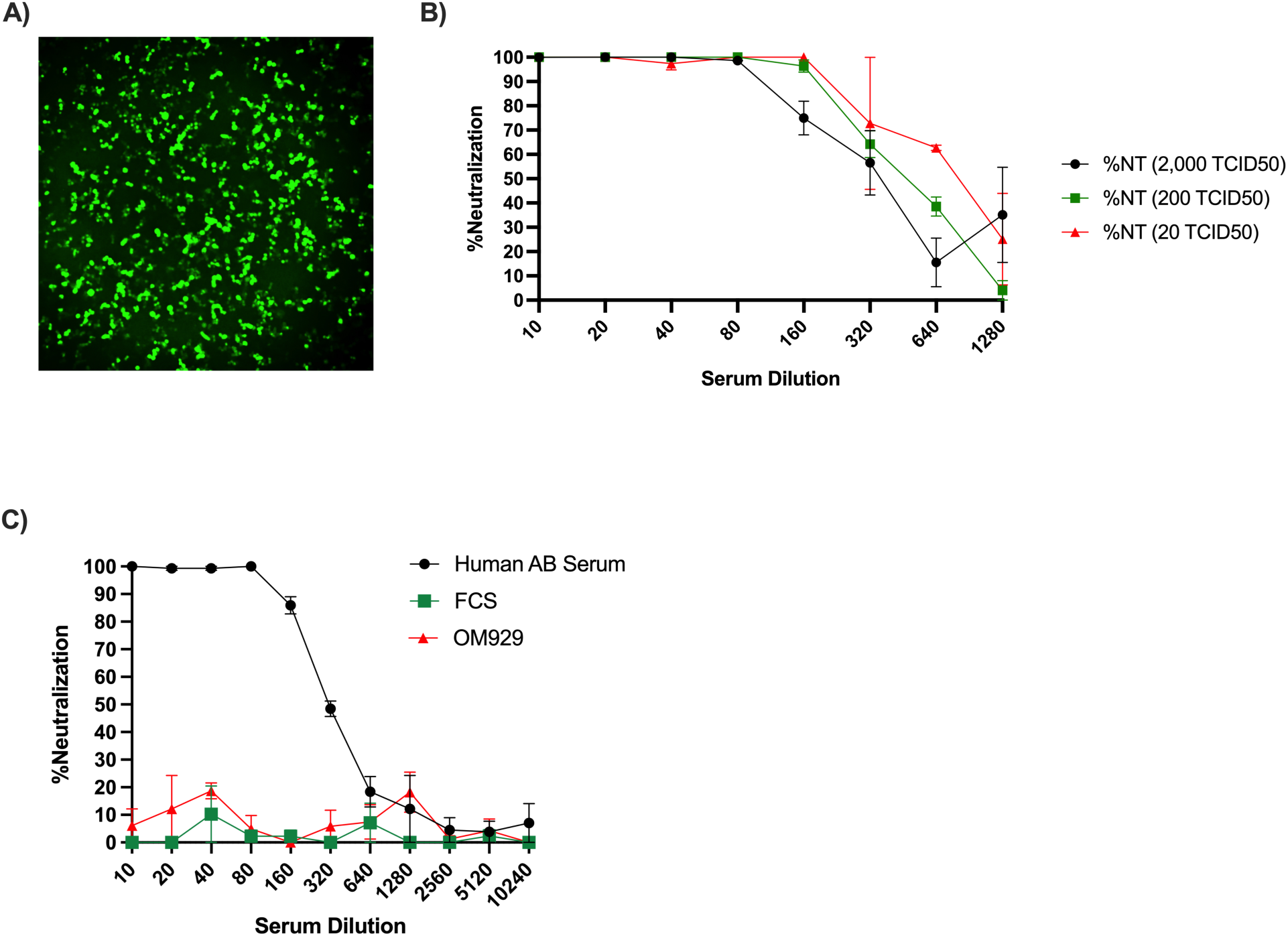
(A) A fluorescence microscopy image of the Akata B-cell line at 48-hours post-inoculation with 17,800 TCID_50_ of EBV-GFP. Green fluorescence confirms successful infection. (B) The EBV-neutralizing capacity of human AB serum across a 10x serial dilution of EBV-GFP (2,000 to 20 TCID50). (C) The serological neutralizing capacity of human AB serum (positive control, black circles), FCS (negative control, green squares), and OM929 (negative control, red triangles). Points and whiskers represent mean values (n = 2 technical replicates) with standard error.

Considering that >90% of individuals are EBV-positive (15), we tested our assay using human AB serum as a positive control. Inoculating the Akata B-cell line with 200 TCID_50_ of EBV-GFP for 48-hours was found to be adequate for measuring serological neutralizing capacity against live EBV (**Figure 4B**). The antibody-dependent neutralizing activity of human AB serum was validated using fetal calf serum (FCS) and an EBV-negative participant (OM929) as negative controls (**Figure 4C**).

Using our assay, live EBV neutralization was compared between anti-VCA IgG seropositive neuropsychiatric long COVID and recovered participants. As shown in **Figure 5A**, EBV neutralizing capacity did not differ between the two groups at 3-6 months PSO. Evidence of EBV reactivation, reflected by elevated EA-D IgG antibody levels, could also only be detected in one neuropsychiatric long COVID participant (**Figure 5B**). However, reduced EBV neutralization at 3-6 months PSO significantly associated with reports of fatigue at 12 months PSO (median EBV IC50 [Fatigue] = 205.40, IQR = 334.38 vs. median EBV IC50 [No Fatigue] = 381.40, IQR = 701.9; p = 0.0435; **Figure 5C**). Anti-VCA IgG antibody levels also strongly correlated with EBV neutralization (r = 0.5706, p < 0.0001; **Figure 5D**) and were significantly diminished in long COVID compared to recovered individuals at 3-6 months PSO (LC median = 149.7 U/mL, IQR = 225.7 vs. R median = 402.1 U/mL, IQR = 314.4; p = 0.026; **Figure 5E**). Anti-VCA IgG antibody levels at 3-6 months PSO similarly trended lower in participants who later reported fatigue at 12 months PSO (median anti-VCA IgG [Fatigue] =149.70 U/mL, IQR = 166.70 vs. median anti-VCA IgG [No Fatigue] = 295.90 U/mL, IQR = 394.10; p = 0.0693; **Figure 5F**).

**Figure 5:**
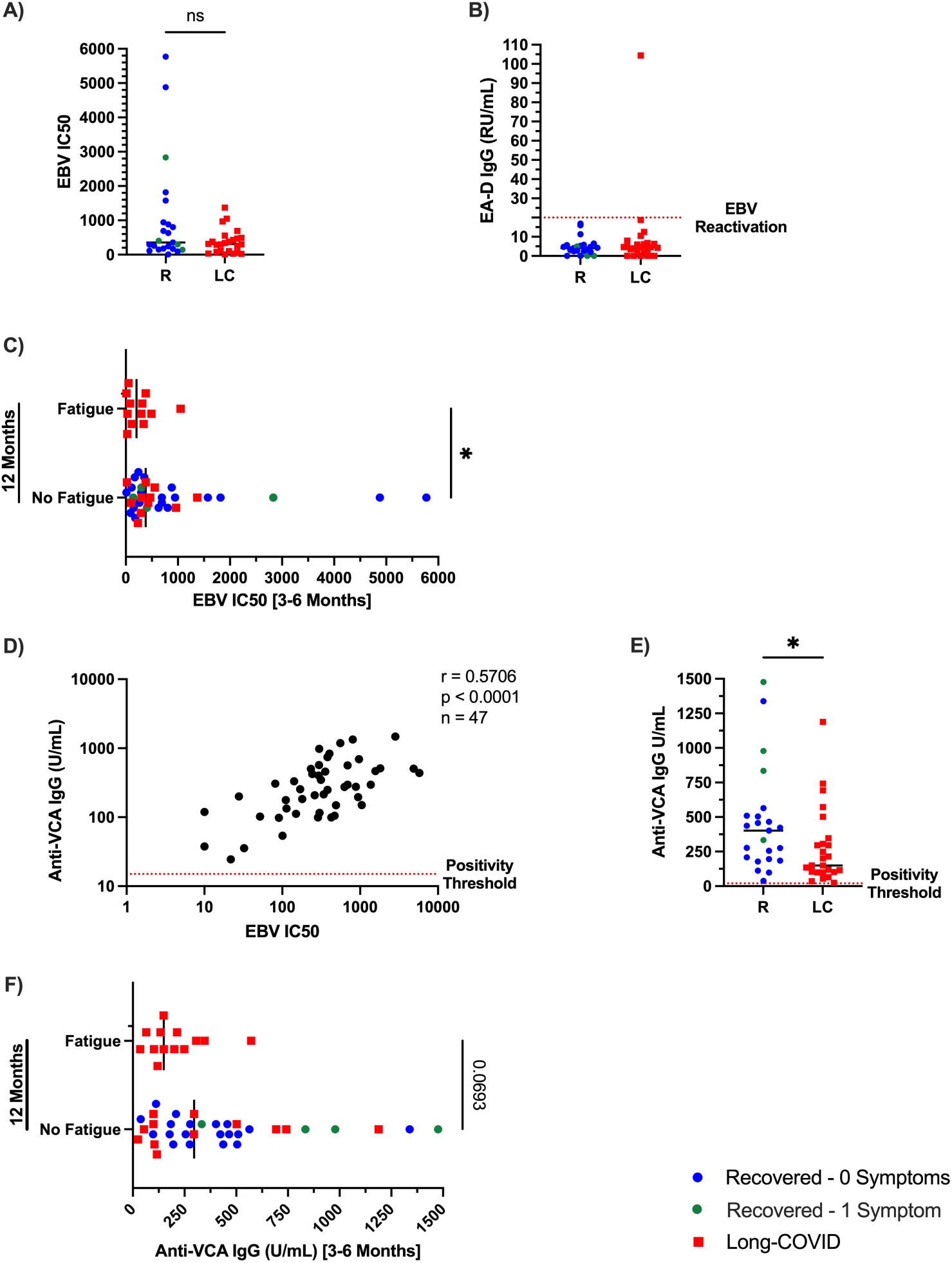
(A) A comparison of EBV neutralizing antibody levels between neuropsychiatric long COVID and recovered participants. (B) A comparison of EA-D IgG antibody titers in relative units (RU)/mL between neuropsychiatric long COVID and recovered participants. The broken red line represents a positivity threshold of 20 RU/mL. (C) A comparison of EBV neutralizing capacity at 3-6 months PSO between participants who experienced no fatigue or fatigue at 12 months PSO. (D) The correlation between EBV neutralizing capacity and anti-VCA IgG antibody levels at 3-6 months PSO. The broken red line represents an anti-VCA IgG positivity threshold of 15 units (U)/mL. (E) A comparison of anti-VCA IgG antibody levels between neuropsychiatric long COVID and recovered participants at 3-6 months PSO. (F) A comparison of anti-VCA IgG antibody levels at 3-6 months PSO between participants who experienced fatigue or no fatigue at 12 months PSO. All solid horizontal lines represent median values.

## DISCUSSION

Long COVID is a multisystemic condition estimated to affect approximately 400 million people worldwide (3, 4). Although underlying mechanisms remain elusive, aberrant ongoing inflammation due to EBV reactivation and persistent SARS-CoV-2 viral reservoirs have been investigated (5, 7–12, 14, 37). A negative correlation between HCMV seropositivity and long COVID incidence has also been reported (41). Here, PBMC and serum samples were collected from 27 long COVID and 27 matched recovered participants at 3-6 months PSO. Four-colour FluoroSpot assays quantified IFN-γ, IL-2, TNF, and GzmB T-cell responses against SARS-CoV-2, EBV, and HCMV. Live viral neutralization assays were additionally conducted to assess serological neutralizing capacity against SARS-CoV-2 and EBV. ELISA assays measuring levels of anti-EA-D IgG, anti-VCA IgG, and anti-HCMV IgG were lastly deployed to confirm EBV reactivation and seropositivity for EBV and HCMV, respectively.

Using four-colour FluoroSpot assays, we could not detect differences in virus-specific T-cell responses between neuropsychiatric long COVID and recovered participants. These findings strongly agree with a study conducted by Williams et al. that likewise compared SARS-CoV-2-, EBV, and HCMV-specific T-cell responses between 22 participants with neurological manifestations of post-acute sequelae of COVID-19 (PASC-N) and 14 asymptomatic recovered participants assessed between 101 and 869 days post-infection (48). Using activation induced marker (AIM) assays and intracellular cytokine staining (ICS) assays for IFN-γ, IL-2, TNF, and GzmB, their study similarly found no differences in T-cell activation or cytokine production in either CD4^+^ or CD8^+^ T-cell subsets following a 24-hour stimulation with SARS-CoV-2-, EBV-, or HCMV-specific peptide pools. As such, it is evident that perturbations in antigen-specific T-cell functionality are unlikely to underlie neuropsychiatric long COVID incidence in either our cohort or that of Williams et al. However, given the modest sample sizes in both studies, larger cohorts examined at similar timepoints post-infection would be needed to validate these findings.

Our live ancestral SARS-CoV-2 neutralization assays also did not uncover any differences in serological neutralizing capacity between neuropsychiatric long COVID and recovered participants. This again withdraws support from our hypothesis and deviates from prior studies conducted by Garcia-Abellán et al. suggesting diminished SARS-CoV-2 neutralization in long COVID individuals (23, 24). However, this is likely attributable to differences in our study cohort: while Garcia-Abellán et al. only reported on long COVID participants who were previously hospitalized, our cohort consisted of those who experienced only mild acute infections. Our findings also agree with Buck et al., who likewise did not detect any associations between neutralizing antibody responses to the ancestral strain of SARS-CoV-2 and long COVID (22). Unlike the previous studies, the cohort examined by Buck et al. was largely non-hospitalized. A limitation of our study, however, is that participant sera were only tested against ancestral SARS-CoV-2, despite infections in our cohort spanning variants from the ancestral strain to Omicron BA.1. This may have potentially masked true differences in neutralization between long COVID and recovered participants. Otherwise, it is not evident that perturbations in SARS-CoV-2-specific neutralizing humoral immunity were associated with neuropsychiatric long COVID incidence in our cohort.

Lastly, among EBV-infected participants, neuropsychiatric long COVID individuals did not exhibit differences in their capacity to neutralize live EBV compared with recovered individuals. However, participants who reported fatigue at 12 months PSO exhibited significantly weaker EBV neutralization at 3-6 months PSO than those who reported no fatigue. This suggests that among EBV-positive participants, weaker EBV neutralizing immunity may have been a risk factor for sustained disease. These observations are supported by a prior report demonstrating that detectable levels of EBV viremia during an acute SARS-CoV-2 infection anticipate sustained disease at 2-3 months PSO (49).

Notably, anti-VCA IgG antibody levels were observed to strongly correlate with EBV neutralization and were significantly diminished in neuropsychiatric long COVID participants at 3-6 months PSO. Diminished levels of anti-VCA IgG at 3-6 months PSO additionally trended with reports of fatigue at 12 months PSO. This finding agrees with a previous report of diminished or absent levels of VCA-antibody secreting cells in individuals with chronic fatigue syndrome (CFS) (28). However, it also deviates from the findings of a more recent report by Klein et al. suggesting that anti-VCA IgG antibodies are elevated in long COVID individuals (50). This may be due to our specific focus on neuropsychiatric long COVID participants compared to a more general long COVID cohort assessed by Klein et al. Deviations in the timing of sampling (3-6 months PSO in our study versus >6 weeks in Klein et al.) may have also contributed to differences in our findings. Nonetheless, our observation of diminished anti-VCA IgG antibody levels potentially suggests a shared mechanism of EBV-specific B-cell dysfunction underlying both CFS and neuropsychiatric long COVID.

Contrary to our hypothesis, through our measurement of EA-D IgG antibody titers, we could only detect 1 incident of EBV reactivation in our long COVID group at 3-6 months PSO (**Figure 5B**). As such, EBV reactivation at 3-6 months was not observed to be associated with long COVID development. However, since we specifically selected participants who contracted a mild SARS-CoV-2 infection, our findings agree with a previous study that likewise could not detect an association between EBV reactivation and long COVID in individuals who contracted asymptomatic or mild acute infections (51).

Another prospective study that investigated the association between EBV reactivation and long COVID development reported a significantly greater incidence of EBV DNA in the throat washings of mildly infected long COVID participants experiencing fatigue compared with asymptomatic recovered individuals (50% vs 20%; p = 0.0411) (14). However, EBV viral load could not be detected within the blood or stool samples of either group. This indicates that EBV reactivation may only be detected within specific tissues following a mild SARS-CoV-2 infection and may not necessarily result in any detectable increases in EBV-specific antibodies including EA-D IgG.

Although Gold et al. observed detectable levels of EA-D IgG in 66.7% of long COVID participants (vs. 10% of recovered individuals) that significantly correlated with the number of symptoms experienced, the study provided little information on the severity of the initial SARS-CoV-2 infection (12). Given several previous reports of a direct positive correlation between EBV reactivation and SARS-CoV-2 disease severity and symptoms (52), EA-D IgG antibodies may be more detectable in long COVID individuals who initially exhibited more severe acute illness.

Within a cohort of 258 long COVID participants, Peluso et al. notably observed that those with detectable levels of EA-D IgG had significantly greater odds of reporting fatigue at a median of 4 months post-symptom onset (53). As per Figure 1 of their study, this timepoint was noted as the general period during which EA-D IgG antibody titers were expected to peak following EBV reactivation (53). Thus, given that the majority of our participants were sampled at 6 months PSO, it is additionally possible that the optimal time to measure EA-D IgG antibody titers was missed. Future studies may need to restrict sampling within a narrower time interval at 3-4 months post-acute symptom onset to minimize variability in the association between EBV reactivation and neuropsychiatric long COVID development.

In summary, our study highlights that neuropsychiatric long COVID symptoms may not be attributable to alterations in SARS-CoV-2-specific cellular or humoral immunity. However, diminished EBV-specific humoral immunity emerged as a potential risk factor for neuropsychiatric long COVID development up to 12 months PSO. Together, these findings may help refine prognostic markers for neuropsychiatric long COVID and inform future mechanistic studies.

## ACKNOWLEDGEMENTS

Live SARS-CoV-2 neutralization assays were conducted by Dr. Patrick Budylowski. Live SARS-CoV-2 virus was provided by Dr. Samira Mubaraka. Hone-Akata cell lines containing EBV-GFP and EBV-negative Akata B-cell lines were generously provided by Dr. Lori Frappier. Victoria Russell and Jeevitha Srighanthan assisted with sample coordination and curated all clinical data used for this study. Dr. Feng Yun Yue managed laboratory operations.

Dr. Angela M. Cheung, who is partially funded by a Tier 1 Canada Research Chair, co-founded and managed the non-ICU portion of the CANCOV research platform for our long COVID studies. This platform was initially funded by the University of Toronto and subsequently funded by CIHR (CANCOV# 172729 and CANCOV 2.0# 179429).

Dr. Mario Ostrowski, supported by the Ontario HIV Treatment Network (OHTN) and the Li Ka Shing Knowledge Institute, provided research direction and expertise throughout the study.

This research was funded by CIHR (20011679 and 20007270) and the Juan and Stefania Fund for COVID-19 and other virus infections.

